# Comparative analysis of two NGS platforms and different databases for analysis of AMR genes

**DOI:** 10.1101/2021.12.27.474239

**Authors:** Twinkle Soni, Ramesh Pandit, Damer Blake, Chaitanya Joshi, Madhvi Joshi

**Author notes:** **Correspondence:** Dr. Madhvi Joshi.

## Abstract

The use of antibiotics in human medicine and livestock production has contributed to the widespread occurrence of antimicrobial resistance (AMR). Recognizing the relevance of AMR to human and livestock health, it is important to assess the occurrence of genetic determinants of resistance in medical, veterinary, and public health settings in order to understand risks of transmission and treatment failure. Advances in Next Generation Sequencing (NGS) technologies have had a significant impact on research in microbial genetics and microbiome analyses. Now, strategies for high throughput sequencing from panels of PCR amplicons representing known AMR genes offer opportunities for targeted characterization of complex microbial populations. Aim of the present study was to compare the Illumina MiSeq and Ion Torrent S5 Plus sequencing platforms for use with the Ion AmpliSeq^™^ AMR Research Panel in a veterinary/public health setting. All samples were processed in parallel for the two sequencing technologies, subsequently following a common bioinformatics workflow to define the occurrence and abundance of AMR gene sequences. Regardless of sequencing platform, the results were closely comparable with minor differences. The Comprehensive Antibiotic Resistance Database (CARD), QIAGEN Microbial Insight - Antimicrobial Resistance (QMI-AR), Antimicrobial resistance database (AR), and CARD-CLC databases were compared for analysis, with the most genes identified using CARD. Drawing on these results we describe an end-to-end workflow for AMR gene analysis using NGS.

## 1. INTRODUCTION

Antimicrobial resistance (AMR) is a growing challenge to the efficient control of diseases caused by bacteria, parasites, viruses, and fungi, prompting the World Health Organization (WHO) to rank it in the top ten public health hazards worldwide^1^. The consequences of AMR include reduced treatment efficacy and increased pathogen persistence, enhancing the likelihood of disease and transmission to others. Multiple-drug-resistant bacteria may already be responsible for 700,000 or more human deaths each year^2^. A report from the UK Antimicrobial resistance review stated that “Advances in genetics, genomics and computer science will likely change the way that infections and new types of resistance are diagnosed, detected and reported worldwide, so that we can fight back faster when bacteria evolve to resist drugs”^3^. One key advance is the use of next generation sequencing (NGS) to detect and analyze the presence of genes and organisms responsible for AMR ^4^.

As sequencing platforms and data analysis pipelines evolve it is important to regularly review their performance in specific applications. An increasingly wide range of NGS platforms are now available, including pyrosequencing, semiconductor based, sequencing by synthesis, and sequencing by ligation ^5^, each based upon a distinct sequencing chemistry. For example, the Ion Torrent technique detects hydrogen ions released during the integration of additional nucleotides into the expanding DNA template^6,7^, while Illumina works on sequencing by synthesis chemistry^8^. Platforms such as Ion Torrent and Illumina also have their own specifications for data quality, read length, total data output per run and library preparation method. Consequentially, it can be challenging for researchers to select the optimal sequencing platform and specifications. The choice of analytical pipeline to process output data adds additional variables, influenced by the nature of the data and the purpose of the study. Several tools are available to detect AMR genes in NGS data, including multiple pipelines, thresholds and databases, hindering comparison between studies. The core objectives of the work presented in this paper were to understand how the use of Illumina or Ion Torrent sequencing platforms impact on data generation, analysis, and final outcome for AMR gene detection in biological samples with relevance to public health.

## 2. MATERIALS AND METHODS

### 2.1 Ethical approval

The work described here was carried out using welfare standards consistent with those established under the Animals (Scientific Procedures) Act 1986, an Act of Parliament of the United Kingdom. All protocols were approved by the Anand Agricultural University (AAU, Gujarat, India) Animal Ethics Committee and the Animal Welfare and Ethical Review Body (AWERB) of the Royal Veterinary College.

### 2.2 Sample Collection and Processing

Twelve apparently healthy broiler chickens (Cobb 400) were collected from the Central Poultry Research Station of Anand Agricultural University, Anand, Gujarat for sampling. All 12 chickens were euthanized by cervical dislocation at 37 days of age. The chickens were reared in a deep litter system using rice husk as substrate, in common with local practices. All chickens were fed a standard maize and soybean-based commercial diet which included bacitracin methylene disalicylate (BMD) and maduramycin (10%) for routine prophylaxis. Samples were collected in RNAprotect Bacteria Reagent (QIAGEN, Germany) as described previously^9^ and transported to the laboratory at 4°C. Upon receipt, total genomic DNA was extracted from each sample immediately using a QIAamp^®^ DNA Stool Mini Kit (QIAGEN, Germany) as described previously^9^. Extracted DNA was stored at −20°C prior to further processing.

### 2.3 AMR Gene Sequencing

While comparing two different sequencing platforms, there should be no difference in the workflow including library preparation. Therefore, we used an Ion AmpliSeq^™^ Antimicrobial resistance (AMR) Research Panel (Thermo Fisher Scientific, MA, USA) for library preparation. This AMR panel consisted of two primer pools targeting 408 and 407 amplicons in each pool. The library preparation flow was also standardized for both the platforms with the exception that Ion-specific adapters and barcodes were ligated for the Ion Torrent platform, while Illumina-specific adapters and indices were used for the Illumina library.

#### 2.3.1 Ion Torrent Platform

Amplicon libraries were prepared using an Ion AmpliSeq^™^ Library Kit Plus (Cat. No. A35907; Thermo Fisher Scientific, MA, USA). Library quality was assessed using a 2100 Bioanalyzer with a DNA high sensitivity assay kit (Agilent CA, USA). Libraries were quantified using the Ion Library TaqMan^™^ Quantitation kit (Cat. No. 4468802; Thermo Fisher Scientific, MA, USA). Sequencing was performed on an Ion S5 Plus system using 530 chip and 400bp chemistry.

#### 2.3.2 Illumina Platform

Amplicon libraries for the Illumina platform were prepared and checked for quality using the same kit and protocol as described above. Sequencing was carried out using an Illumina MiSeq system with a MiSeq reagent kit v2 and 500 cycles (250 x 2 paired end chemistry).

### 2.4 Data Analysis

Data obtained from Ion torrent and Illumina MiSeq was analyzed using the same bioinformatics pipeline. The initial difference in the paired end read from Illumina and single end reads from Ion torrent was nullified by merging the paired end reads of Illumina using PandaSeq v 2.8.1^10^. Here also, different overlapping parameters were first assessed for the best results such as 5bp, 10bp, 15bp and default overlapping.

The quality of raw data was assessed using FastQC v. 0.11.5^11^. The average quality score threshold to retain a read was set 30 for Illumina and 20 for Ion Torrent data because of the inherent differences in the base calling accuracy due to differences in the sequencing chemistry of these two platforms^12,13^. Read trimming for length was not performed as the smallest amplicon targeted in the panel was 72bp. CARD database version 3.0.7^14^ was used for analysis. Local BLASTn or BLASTx was performed with the following parameters: no. of alignments retrieved 1, minimum percent identity 95% and E-value 10e^-5^. The downstream statistical analysis was done using Excel and STAMP v2.1.3^15^.

### 2.5 Comparison of Different Databases

Four different databases namely, Comprehensive Antibiotic Resistance Database (CARD)^14^, QIAGEN microbial Insight – Antimicrobial Resistance (QMI-AR)^16^, Antimicrobial Resistance (AR), and CARD-CLC^17^ were used for comparison. These databases were compared using stringent parameters including number of alignments per read as 1, minimum alignment length as 95%, E-value as 10 e-5 and percent identity for BLAST as 95%. The downstream analysis was performed using STAMP and Venny 2.1.0^18^.

### 2.6 Microbiome Analysis

The Online web-based tool Microbiome Analyst^19,20^ was used to perform LEfSe (Linear discriminant analysis Effect Size), PCoA (Principle coordinate analysis) with PERMANOVA statistics, and Random forest analyses in order to support statistical comparison. In LEfSe, Log LDA cutoff was set as 3.0 with p-value ≤0.05.

## 3. RESULTS

To support national and global priority setting, public health initiatives, and treatment decisions, a credible base of knowledge that appropriately captures and characterizes the worldwide burden and transmission of AMR is required. In this study, efforts were made to compare different sequencing technologies and data bases to provide a comprehensive analysis pipeline for data defining AMR gene occurrence. All experimental variables were fixed with the exception of sequencing platform. Library preparation kit, data analysis pipeline, and database stringency were all kept the same to maintain uniformity.

### 3.1 Sequencing results

In total ~15M reads were obtained using the Ion Torrent S5 Plus platform in a single FastQ file, representing approximately 1M reads per sample with an average read length of 200bp. In parallel, 4.18M reads were produced for the same samples using Illumina MiSeq, representing 0.2M reads per sample with an average read length of 185 bp.

### 3.2 Optimization of overlapping parameter for Illumina

Based on our previous experience with PandaSeq in merging 16S Illumina amplicon data and the toll’s citations (>1600), we selected this toll for this study. Classification of Illumina forward and reverse reads required attention because of issues when merging paired end reads^21^. Initially, we merged reads using PandaSeq’s default parameters; later, the merge length was optimized. Forward-reverse read overlap of 5, 10, and 15bp was analyzed in addition to the default parameters. The 10 base pair overlap was found to be optimal due to its appropriate representation of merged reads (Fig: S1). These results showed that overlapping parameters for merging forward-reverse amplicon reads may incur important differences in apparent gene abundance as an appropriate overlapping parameter leads to false positive and negative results in Illumina sequencing platforms while analyzing AMR data.

### 3.3 Optimization of BLAST parameters

The BLAST (Basic Local Alignment Search Tool) algorithm is used widely, but output is influenced by the parameters applied. Therefore, in this study, various BLAST parameters were optimized along with the overlapping length used in PandaSeq. Three conditions were set, being default BLAST and default overlap, 10bp overlap and default BLAST and 10bp overlap and BLAST query hsp percentage 90 (Fig:S2). The default overlap with default BLAST could not be used for analysis due to nonspecific reads merges. Specifically, the PandaSeq default merge length is 1bp, indicating that any two reads possessing a common base at the 5’ will be merged. The 10bp overlap and BLAST qcov hsp percentage 90 was also not efficient as it hampered estimation of occurrence for genes such as *ErmB*. The 10bp overlap with default BLAST was found to be most accurate as it avoided these issues and was applied for all subsequent analyses.

### 3.4 Comparison of Ion Torrent and Illumina MiSeq for AMR gene detection

More AMR genes were detected using the Ion Torrent Platform compared to Illumina MiSeq (average number of genes detected 369±58 compared to 206±38, respectively from all 12 samples). In total, the Ion Torrent platform detected the presence of 31.9% more AMR relevant genes compared to Illumina MiSeq, although the percentage abundance of these genes was very low (i.e. less than 0.004%). Additionally, 6% of genes detected using Illumina MiSeq were missing from the Ion Torrent results, but again the percentage abundance of each gene found only by Illumina was very low (i.e. less than 0.004%). There were many genes that were only discovered in one or two samples, and those with a small number of hits. Overall, 62.1% genes detected were common across both platforms. But, when genes with abundance ≥1% sequencing reads were considered, the results from both sequencing platforms were similar (Table:1, Fig:1). The APH (3’)-IIIa gene was found to be most abundant in both the platforms followed by *tetW* and *tetQ*. The occurrence of only nine genes was found to be significantly different between the sequencing platforms (Fig: S3). Out of these nine genes, *tet(40)* was found to be most variable (4%). Sample-specific comparison highlighted similar platform-associated variation for the occurrence of *tetO* and Aminoglycoside phosphotransferase genes (Fig: S4, Fig: S5). Direct sample-specific comparison revealed comparable gene detection profiles using Illumina MiSeq and Ion Torrent S5 Plus for genes with greater than 1% read abundance (Fig:2).

**Table:1.** Comparative analysis of the presence or absence of AMR genes represented by ≥1% sequence abundance within Illumina MiSeq or Ion Torrent amplicon sequencing datasets.

|  | **Number of AMR genes detected** |  |  |  |
| --- | --- | --- | --- | --- |
| **Sample** | **Total** | **Illumina AND**<br>**Ion Torrent** | **Illumina** | **Ion Torrent** |
| S1 | 19 | 18 | 1 | 0 |
| S2 | 16 | 15 | 1 | 0 |
| S3 | 13 | 13 | 0 | 0 |
| S4 | 14 | 14 | 0 | 0 |
| S5 | 13 | 13 | 0 | 0 |
| S6 | 12 | 12 | 0 | 0 |
| S7 | 13 | 13 | 0 | 0 |
| S8 | 16 | 15 | 1 | 0 |
| S9 | 18 | 14 | 1 | 3 |
| S10 | 16 | 11 | 0 | 5 |
| S11 | 16 | 12 | 3 | 1 |
| S12 | 16 | 14 | 2 | 0 |

**Figure:1.**
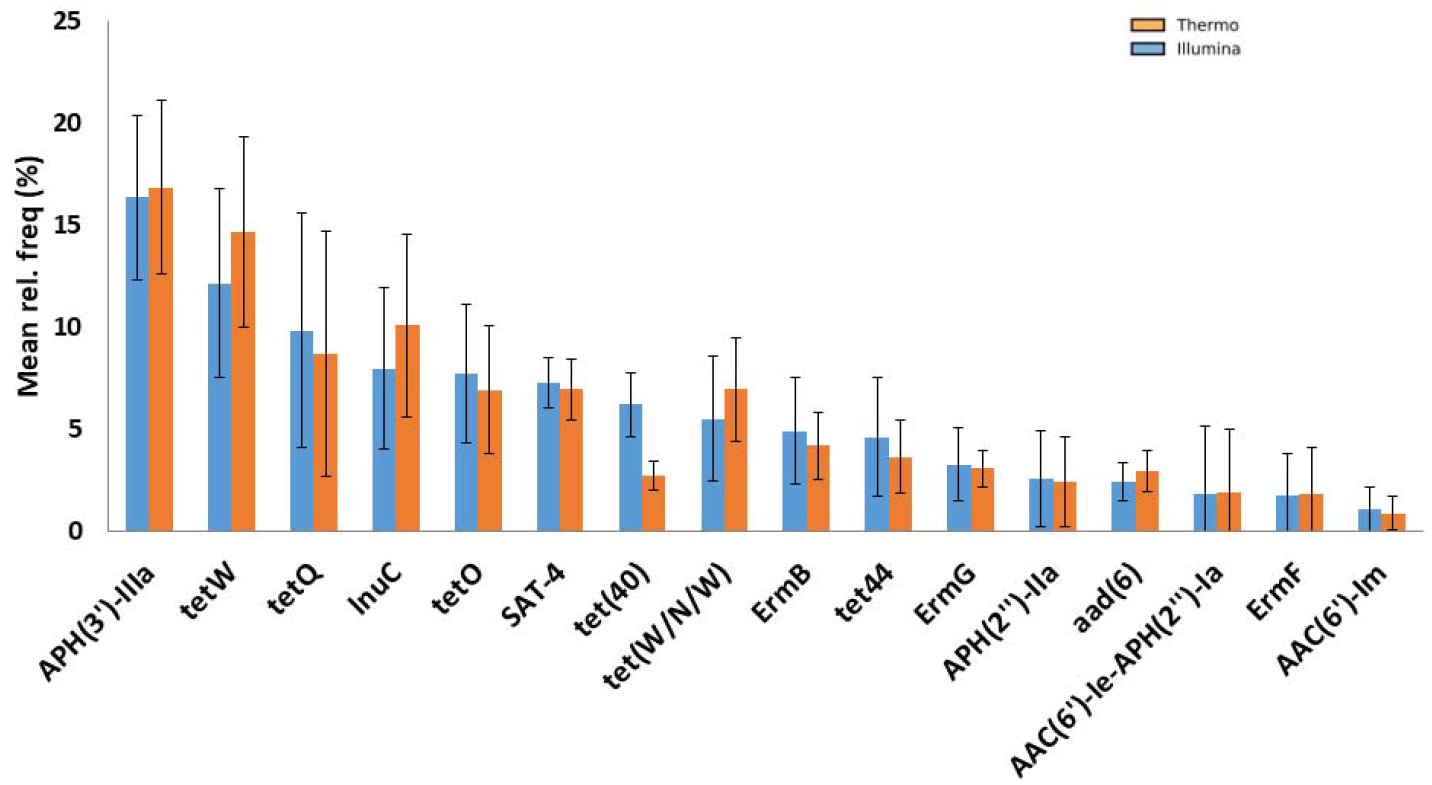
The relative sequencing read abundance of AMR genes amplified from chicken caecal microbial populations with ≥1% abundance within equivalent Illumina MiSeq and Thermo Fischer Scientific

**Figure:2.**
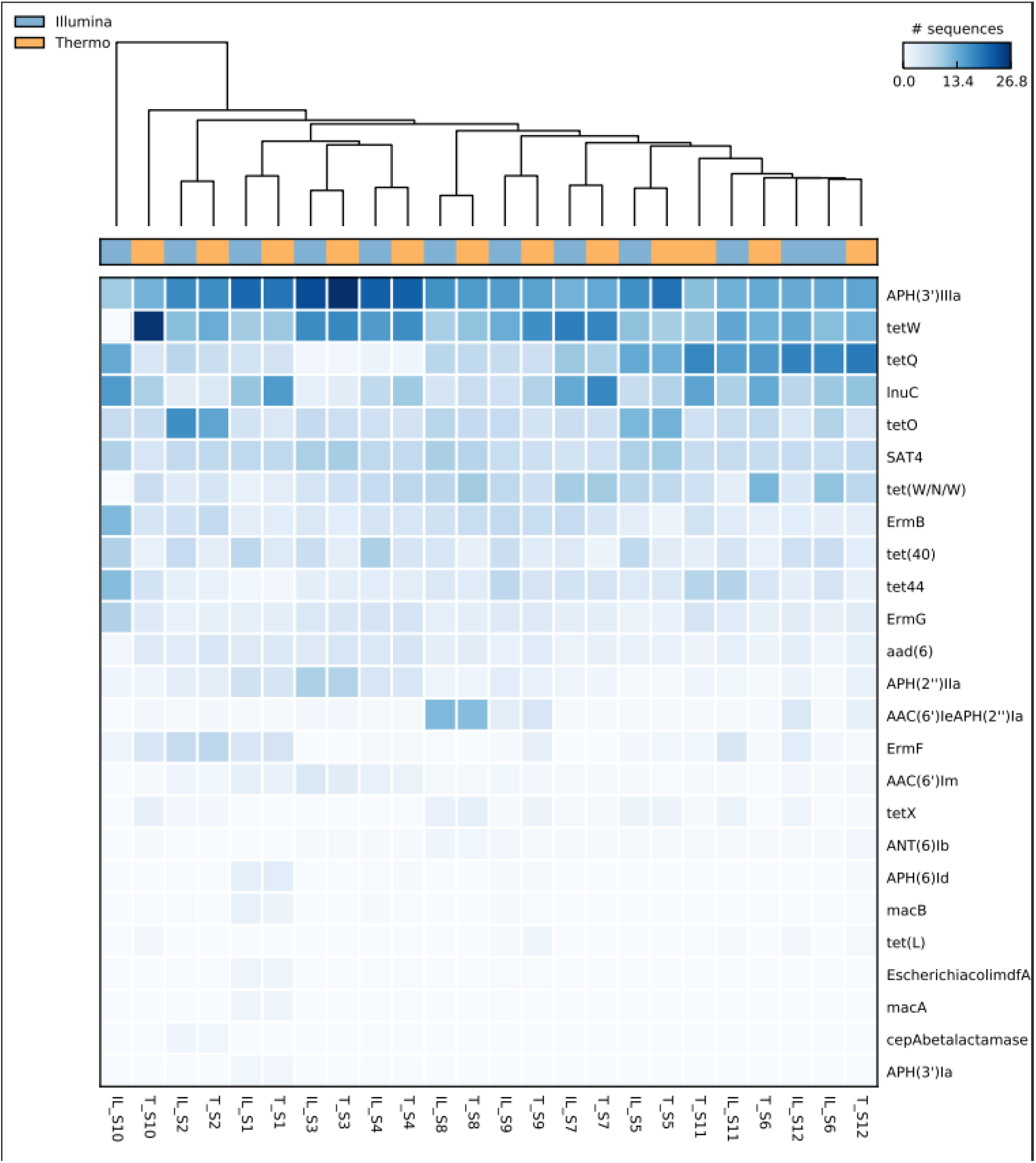
Heatmap demonstrating the abundance of the top 25 AMR genes in 12 chicken caecal samples detected using Illumina MiSeq (IL) or Ion Torrent (T), plotted using STEMP. AMR gene abundance illustrated by a color gradient as indicated by the color scale.

### 3.5 *Tet(40)* and Lnu C Comparison

The abundance of *tet(40)* was found to be higher using the Illumina MiSeq platform (6.21±1.26 %) when compared to Ion Torrent (2.5±1.0%). Annotation using the CARD database indicated *tet(40)* carriage by a group of uncultured bacteria. Thus, a comparable trend was observed when samples were compared by predicted contributing organism (Illumina 6.2±1.6%; Ion Torrent 2.5±1.0%). In contrast, *lnuC* was more highly abundant in the Ion Torrent dataset (9.79±5.15%) compared to Illumina (7.9±4.1%) (Table: 1). The *lnuC* gene was predicted to be carriedby *Streptococcus agalactiae* and hence, the same trend in the percentage of *S. agalactiae* could be observed (Fig:1, Fig:3).

**Figure:3.**
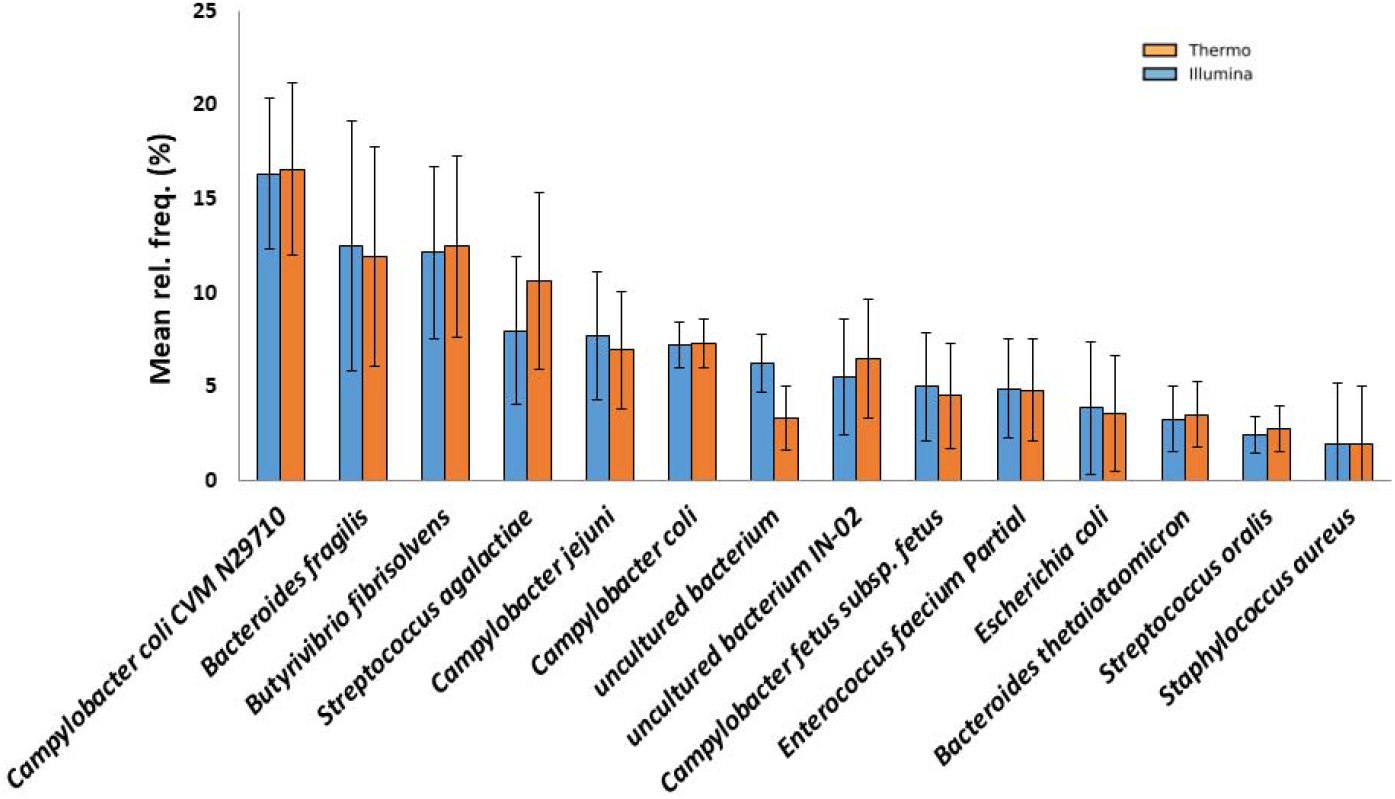
The relative abundance of organisms hosting AMR genes in chicken caecal microbial populations predicted using the CARD database. Organisms represented by ≥1% of the sequence reads generated using Illumina MiSeq or Thermo Fischer Scientific are shown.

### 3.6 Microbial diversity comparison on the basis of AMR detected

Prediction of bacterial identity associated with AMR gene carriage was found to be comparable in both the platforms (Table:2, Fig:3). *Campylobacter coli* CVM N29710 was the most abundant organism identified, followed by *Bacteroides fragillis*. Only *Staphylococcus epidermidis* was found to be significantly differently represented e between the platforms (q-value (corrected) = 0.001) (Abundance <0.0014) (Fig: S6). Comparison of bacterial representation in individual samples as also undertaken, illustrating the stability of taxonomic classification between sequencing platforms (Fig 4, Fig: S7, Fig: S8).

**Table:2.** Comparative analysis of the presence or absence of organisms predicted to host AMR genes detected within Illumina MiSeq or Ion Torrent amplicon sequencing datasets. Organisms representing AMR genes with ≥1% abundance are shown.

| Sample | Number organisms identified by CARD |  |  |  |
| --- | --- | --- | --- | --- |
|  | Total | Illumina AND<br>Ion Torrent | Illumina | Ion Torrent |
| S1 | 15 | 14 | 0 | 1 |
| S2 | 13 | 13 | 0 | 0 |
| S3 | 12 | 12 | 0 | 0 |
| S4 | 13 | 13 | 0 | 0 |
| S5 | 12 | 12 | 0 | 0 |
| S6 | 13 | 12 | 1 | 0 |
| S7 | 13 | 13 | 0 | 0 |
| S8 | 14 | 14 | 0 | 0 |
| S9 | 15 | 14 | 0 | 1 |
| S10 | 14 | 10 | 0 | 4 |
| S11 | 14 | 13 | 1 | 0 |
| S12 | 14 | 14 | 0 | 0 |

**Figure:4.**
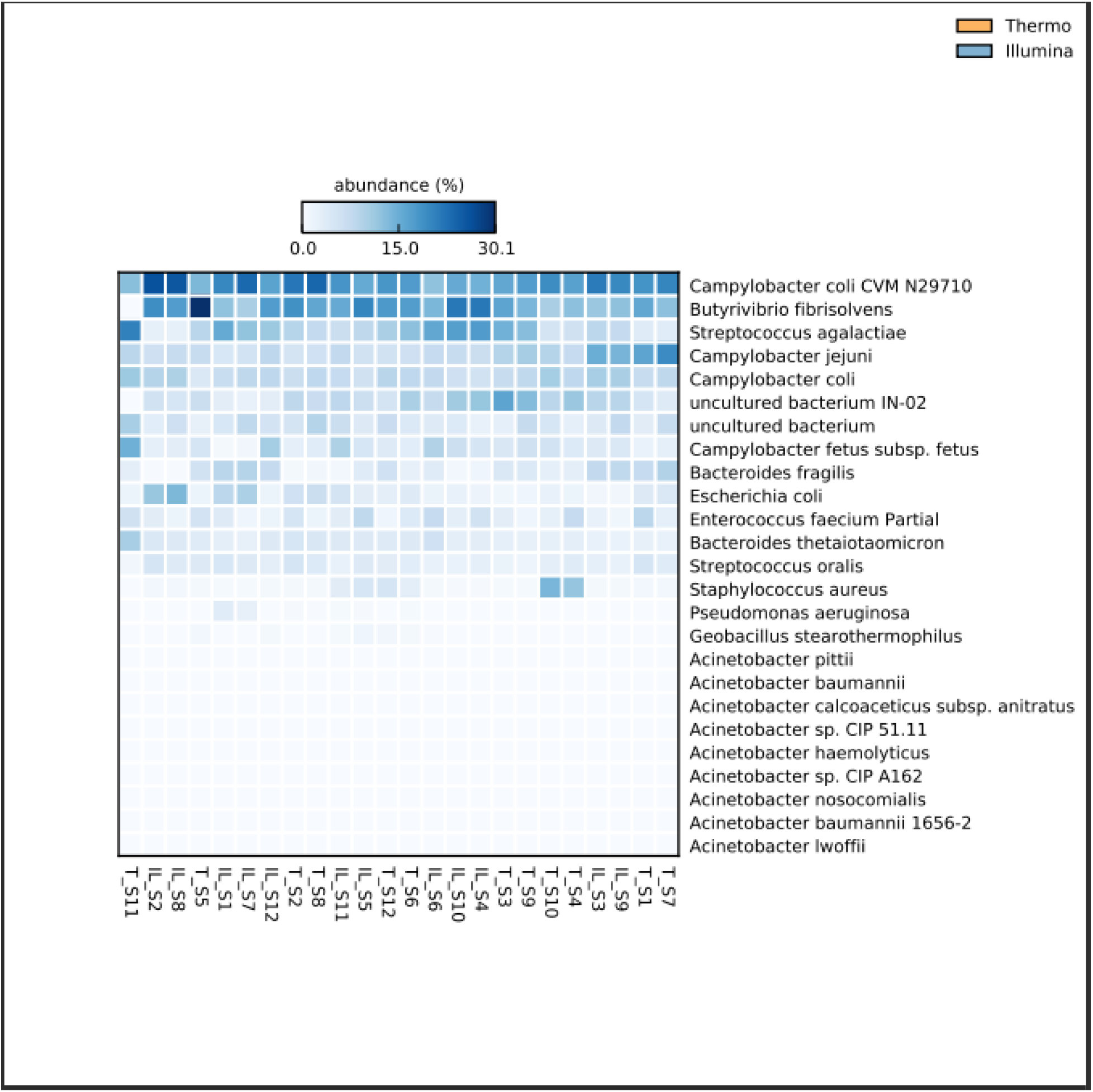
Heatmap demonstrating occurrence of the top 25 most abundant organisms predicted in chicken caecal microbial populations using the CARD database. Data generated from equivalent Illumina MiSeq (IL) and Ion Torrent (T) AMR amplicon sequencing datasets and plotted using STEMP. Organism abundance illustrated by a color gradient as indicated by the color scale.

### 3.7 Database comparison

Several databases are available for analysis of AMR genes. Comparison of the CARD, QMI-DB, AR and CARD_CLC databases with stringent parameters produced varied results with limited correlation or similarity (Fig: 5). In the absence of clear complementarity, the CARD database was chosen for downstream analysis because it is easily available and hosts the largest number of genes and organisms among the four databases. As CARD is used primarily with genome sequence data, a ‘model’ of detection for each sequence means that the criteria that determine the corresponding sequence for each sequence of the CARD reference are determined. The microbiological analysis module in CLC genomic workbench (version 21.1) was utilized to compare results. The investigation also made use of the CARD database in CLC genomic workbench

**Figure:5.**
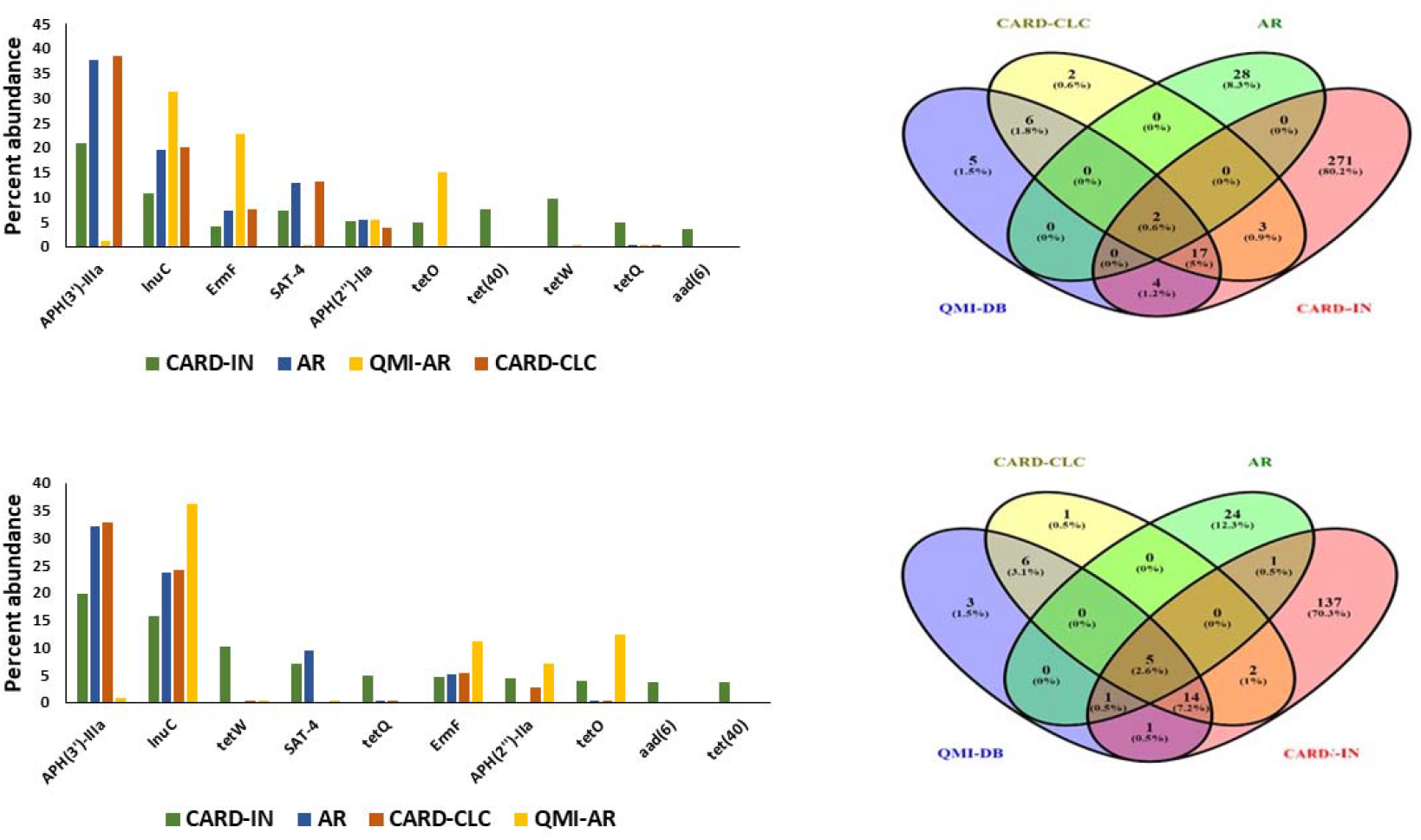
Comparison of sequence analysis databases for assessment of AMR gene amplicon sequencing generated using Illumina MiSeq and Ion torrent. The figures on the top row represts the data from Illumina platform while data of down row represents the Ion Torrent platform. Databases included were (CARD-CLC-CARD database present in CLC genomic workbench microbial genomic module, AR – Antibiotic resistance database, QMI-DB-QIAGEN microbial Insight – AR, CARD-IN – CARD database downloaded from CARD site and run locally)

### 3.8 Statistical comparison of AMR gene occurrence detected by Illumina MiSeq and Ion Torrent sequencing

The random forest method generates decision trees from data samples, generation multiple predictions before identifying the best solution. Random forest is an ensemble method that is superior to a single decision tree because it averages results to reduce over-fitting (Pedregosa et al., 2011). Here, random forest analysis was performed in order to identify any outliers in each dataset. Comparison of Illumina MiSeq and Ion Torrent datasets revealed the absence of outliers, supporting the accuracy of each (Fig: S9). Similarly, PCoA analysis was used to confirm that all Illumina MiSeq and Ion Torrent sample sequences were located in the same cluster (Fig. 6). For AMR gene and organism comparisons there were no significant differences (PERMANOVA; F-value 1.3421, R2 value 0.057498, p-value <0.219; and F-value 0.82178, R2 value 0.036009, p-value <0.514; respectively).

**Figure:6.**
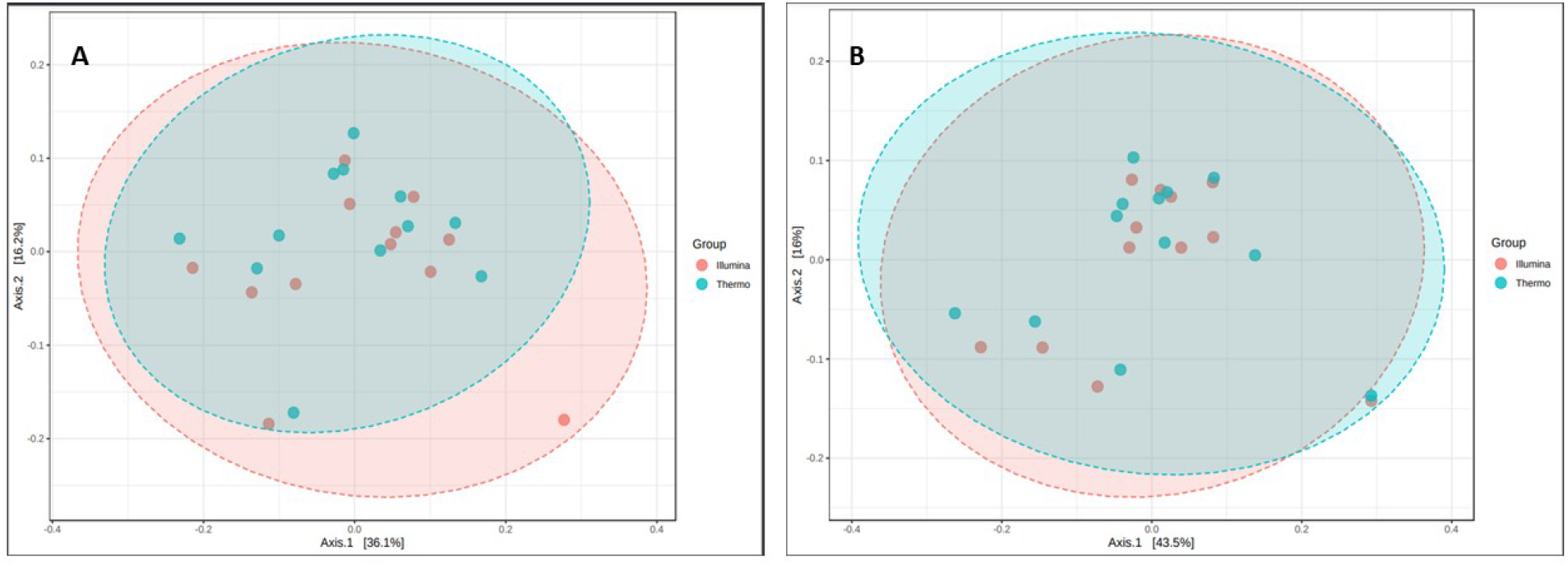
PERMANOVA analysis of AMR gene and organism occurrence predicted based upon Illumina MiSeq or Ion Torrent sequencing of AMR amplicons from chicken caecal contents. (A) AMR gene PERMANOVA with F-value 1.3421, R^2^ value 0.057498 and *p-value <0.219 (B) Organism PERMANOVA analysis with F-value 0.82178, R^2^ value 0.036009 and *p-value <0.514.

### 3.9 LEfSe analysis

LEfSe was performed for both gene and organism at Log LDA 3.0 and P-value ≤0.05. Only four of 300 organisms were found to be significantly different between sequencing platforms. *Enterococcus faecalis*, Plasmid_pGT633 and *Bacteroides coprosuis* was more abundant in the Ion Torrent dataset while, Uncultured bacteria were more common using the Illumina platform. However, the abundance of all four organisms was low, less than 0.07% and 0.02% in the Ion Torrent and Illumina datasets, respectively, both below the 1% threshold set earlier (Fig: S10). LEfSe analysis of the AMR genes detected indicated that five genes were significantly different between the platforms (Fig: S11). The genes *tet32, ErmT, tetS* and *Erm35* was found to be more abundant in Ion Torrent sequencing, while *tet(40)* was more common in the Illumina data. Again, the percent abundance of these gene-specific reads was less than 0.04% in the Ion Torrent sequencing. Only detection of the gene *tet(40)* was found to be significantly different with more than 1% read abundance, presenting with a two-fold higher abundance in the Illumina MiSeq data.

## 4. DISCUSSION

The study was planned to answer the very basic question associated with the use of NGS sequencing platform for AMR analysis. Therefore, in this study we compared the two sequencing platforms, Ion Torrent and Illumina MiSeq for the analysis of AMR and set abioinformatics data analysis pipeline after consideration of all the difference between two platforms. The study was performed with 12 chicken cecum samples to estimate the abundance of AMR genes and corresponding organisms.

The bioinformatics pipeline generated for the data analysis was tried to keep constant for both platforms. Although the initial parameters like quality score threshold and overlapping parameter vary a bit in both the platforms. Due to higher confidence at quality score greater than 30 in Illumina and greater than 20 at Ion torrent we had set different initial quality cutoffs for the data. In addition to this, Ion torrent results in single-end sequencing while paired-end sequencing was used in case of Illumina. In order to merge the forward and reverse reads of illumina an extra step of reads merge was performed. These two changes bring data from both the platform on same page. Later, the local blast parameters and data analysis parameters were kept stringent and constant for both the data sets.

Upon the completion of analysis, it was found that Ion Torrent resulted in higher numbers of hits (31.9%) in case of AMR detection as compare to Illumina MiSeq platform. However, those hits were found to be insignificant as their percent abundance was less than 0.004%. The qualitative and quantitative similarity was found among the significant AMR genes. The similar results were obtained by Lahens et al., upon the comparative analysis of differential expression of gene among ion torrent and illumina^22^. The difference in other insignificant hit may arise from the sequencing errors and poor quality. The *tet*(40) gene was found to be significantly different among both the platforms. Upon detailed analysis of *tet*-(40) abundance, it was found that the amplicon length of *tet*racycline 40 gene is 80bp only. It is among the shortest amplicon present in the AMR panel. This short amplicon length may result in the phenomenon of competitive binding on Illumina flow cell while cluster generation. Competitive binding means shorter amplicon tends to bind to flow cells more as compared to the larger one. The bacterium which corresponds to this *tet(40)* is uncultured bacteria. Hence, the same trend was observed in the uncultured bacteria. Inverse to this, during emulsion PCR of Ion Torrent, the shorter fragments tend to form polyclonal and therefore, the reads tends to be discarded. Therefore, we expected that, this could be one of the possible reasons for *tet(40)* gene’s lesser abundance in Ion Torrent and higher in Illumina dataset. Similarly, Lincosamide resistance gene is one of the largest amplicons in AMR panel (224bp). The phenomenon opposite to that of *Tet-40* may work here i.e. lower abundance of *LnuC* in Illumina data as compared to Ion torrent. The *LunC* gene is mostly contributed from the *S. agalactiae* so a similar trend is observed there as well. The only statistically significant difference of only one organism was found i.e. *Staphylococcus epidermidis.* The variation in the abundance of *S. epidermidis* is almost negligible as its abundance is very less.

Two different platforms were used to identify any database correlation if any. One of these platform was CARD local database and another was CLC workbench with QIAGEN microbial insight module providing different database for the AMR search (QMI-AR, AR, CARD). CARD local database was preferred due to its capacity to target higher number of gene as compare to another. Moreover, the main disadvantage of CLC workbench is that it is not freely available. Both the CLC workbench license and the microbiological insight module have separate costs to pay.

The present study has effectively demonstrated that, the analysis platform used to detect AMR in samples does not significantly influence the results. On analyzing the sample costs and availability of the instrument, the selection of platform is advised. The only limitation of the present study is; we do not perform the same exercise on the mock community as, such mock community was not available.

## 5. CONCLUSION

Irrespective of sequencing chemistry and platform used, comparative analysis among AMR genes and candidate host organism suggest that the Illumina MiSeq and Ion Torrent platforms performed equally. According to the findings, authors suggest that using any platform or sequencing chemistry has little effect on the outcome. In both the platforms APH-IIIa was the most abundant AMR gene. Comparative analysis of the organisms identified in each sample rarely varied significantly. In both the platforms, *C. coli* CVM N29710 was the most abundant bacterium. The statistical significance difference among the t*et*(40) gene was observed which may arise with the short length amplicons. Furthermore, in order to correctly assess AMR in biological samples, standard methods and pipeline for sample analysis must be established. Database selection and parameter for analysis can change the outcome considerably.

**Table:3.** Variation in the relative abundance of *tet(40)* and *lnuC* gene amplicons detected in chicken caecal bacterial populations using Illumina MiSeq or Ion Torrent amplicon sequencing. Likely host organism (as predicted by CARD) and amplicon length is shown.

| GENE | ILLUMINA | ION TORRENT | ORGANISM | AMPLICON<br>LENGTH |
| --- | --- | --- | --- | --- |
| *tet(40)* | High (6.2%±1.3 %) | Low (2.5%±1.0%) | Uncultured<br>bacteria | 80 |
| *lnuC* | Low (7.9%±4.1%) | High (9.8%±5.1%) | *S. agalactiae* | 224 |

## AUTHOR CONTRIBUTIONS

Twinkle Soni: Data curation, Writing - Original draft preparation; Response, Analyses and Illustrations.

Ramesh Pandit: Review & Editing manuscript.

Damer Blake: Review & Editing manuscript.

Madhvi Joshi: Review & Editing, Validation.

Chaitanya Joshi: Supervision and Validation.

## ACKNOWLEDGEMENTS

The authors like to acknowledge One Health Poultry Hub for the support.

## Supplementary figure

**Figure: S1** Optimization of overlapping parameters in PandaSeq while keeping rest parameter same (R1-Forward sequence, R2-Reverse sequence, Default merge-without adding any overlapping criteria manually, 5 base OL-minimum overlap of 5 base pair, 10 base OL-minimum overlap of 10 base pair, 15 bases OL-minimum overlap of 15 base pair).

**Figure: S2** Optimization of blast parameters while keeping rest parameter same (Default overlap; Default blast-No manual parameters added, 10bp overlap; qcov_hsp_perc 90 blast-10 base pair overlap in PandaSeq with query hsp percentage set 90 for blast result, 10bp overlap; Default blast-10 base pair overlap in PandaSeq and default blast parameters).

**Figure: S3** STEMP analysis of total gene found in Illumina and Ion Torrent for estimation of statically significant difference among the abundance of gene in both the platform.

**Figure: S4** Sample vise comparison in abundance of *Tet*racycline-O gene among all the samples on different platforms. (p=0.775).

**Figure: S5** Sample vise comparison in abundance of Aminoglycoside phosphotransferase, gentamicin resistance protein gene among all the samples on different platforms. (p = 0.901).

**Figure: S6** STEMP analysis of total organism found in Illumina and Ion Torrent for estimation of statically significant difference among the abundance of organism in both the platform.

**Figure: S7** Sample vise comparison in abundance of *Campylobacter jejuni* among all the samples on different platforms. (p = 0.799).

**Figure: S8** Sample vise comparison in abundance of *Campylobacter fetus subsp. fetus* among all the samples on different platforms. (p = 0.876).

**Figure: S9** Random forest classification of Illumina vs Ion Torrent samples depicting the confidence in data analysis.

**Figure: S10** The LEfSe analysis of the organism keeping log LDA cut off as 3.0 and p-value as 0.05.

**Figure: S11** The LEfSe analysis of the AMR genes keeping log LDA cut off as 3.0 and p-value as 0.05.

