## Supplementary material for "Comparative analysis of two NGS platforms and different databases for analysis of AMR genes": MD Supplementry.docx

**SUPPLEMENTRY FIGURES**

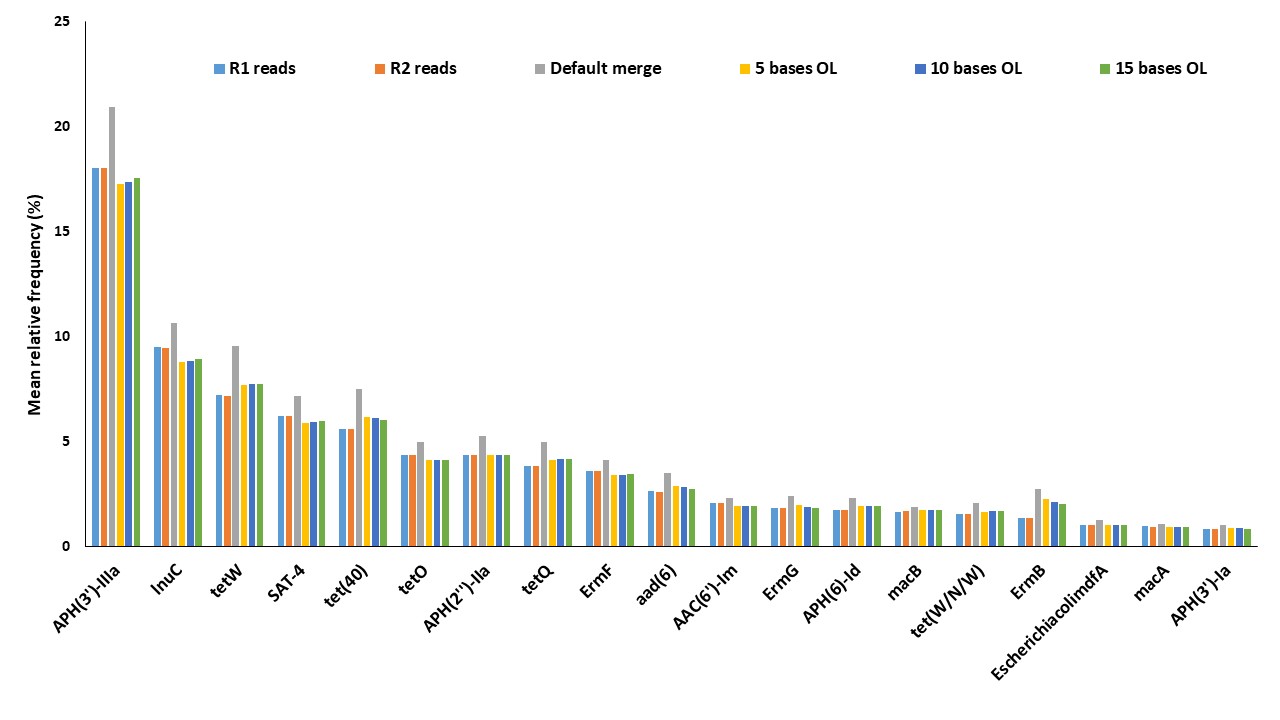

Figure: S1 Optimization of overlapping parameters in PandaSeq when other parameters were fixed (R1- Forward sequence, R2- Reverse sequence, Default merge- without adding any overlapping criteria manually, 5 base OL- minimum overlap of 5 base pair, 10 base OL- minimum overlap of 10 base pair, 15 bases OL- minimum overlap of 15 base pair)

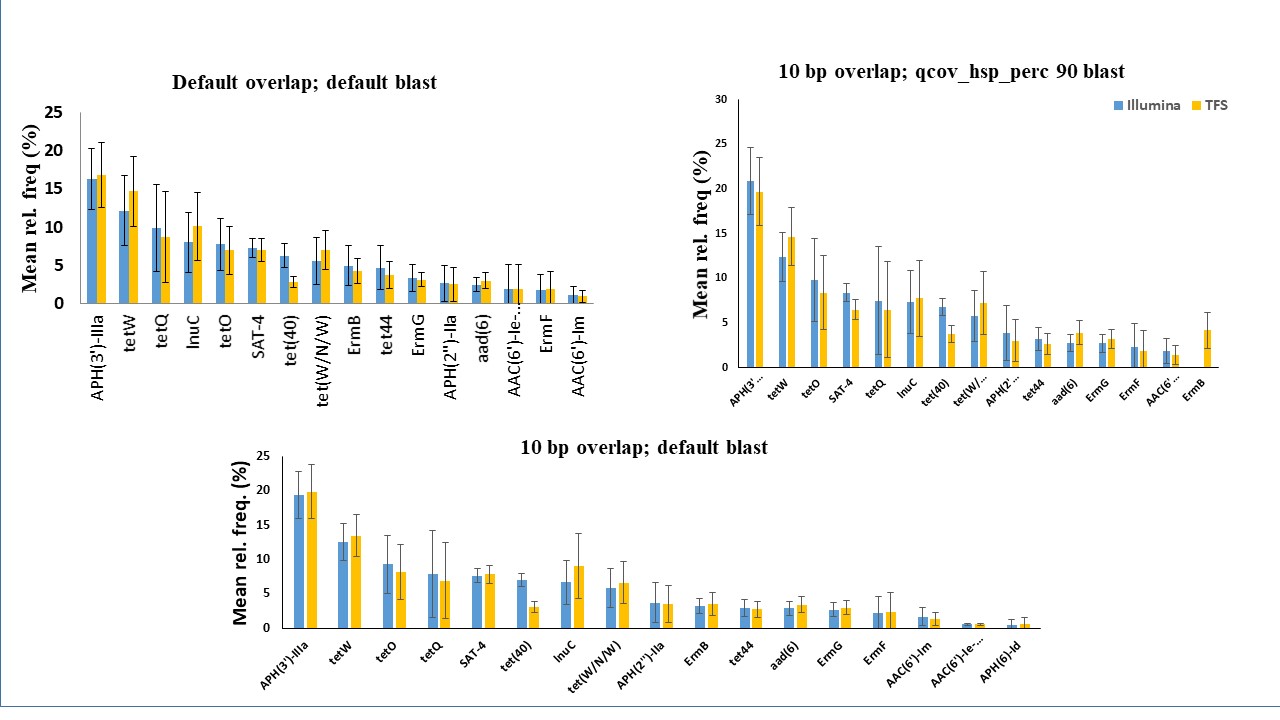

Figure: S2 Optimization of blast parameters when other parameters were fixed (Default overlap; Default blast- No manual parameters added, 10bp overlap; qcov_hsp_perc 90 blast- 10 base pair overlap in PandaSeq with query hsp percentage set 90 for blast result, 10bp overlap; Default blast- 10 base pair overlap in PandaSeq and default blast parameters). Illumina- Illumina MiSeq, TFS- Thermo Fischer Scientific

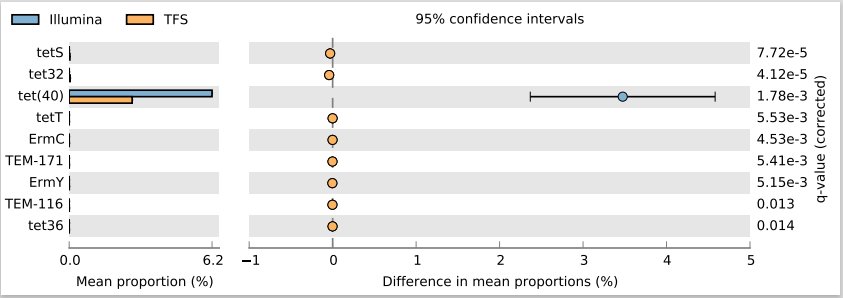

Figure: S3 STEMP analysis of total gene (~885) found in Illumina MiSeq and Ion Torrent datasets to estimate statistically significant differences in gene abundance. Illumina- Illumina MiSeq, TFS- Thermo Fischer Scientific

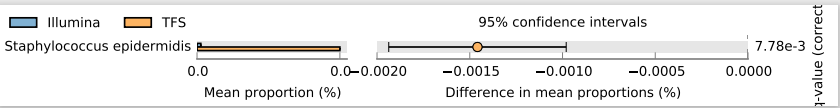

Figure: S4 STEMP analysis of total organism found in Illumina and Ion Torrent for estimation of statically significant difference among the abundance of organism in both the platform

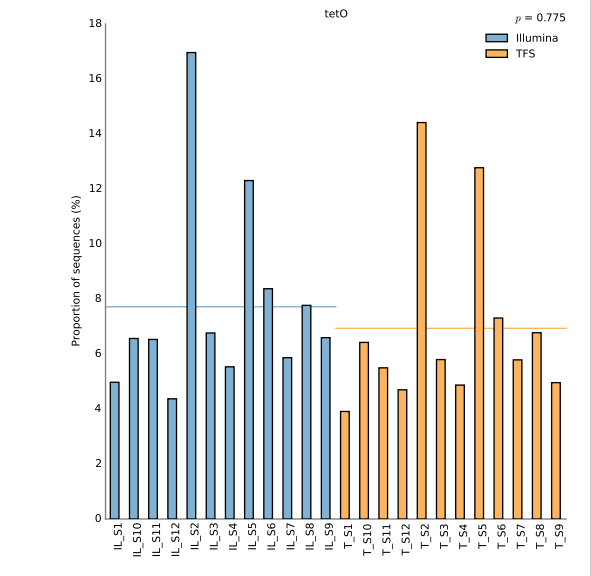

Figure: S5 Comparison of Tetracycline-O gene abundance in individual samples sequenced using Illumina MiSeq or Ion Torrent platforms. (*p=0.775)

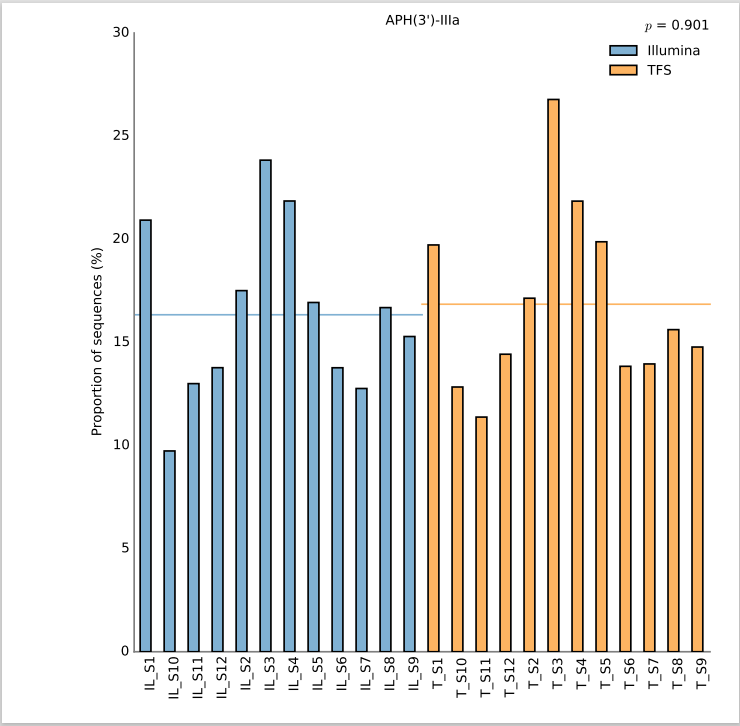

Figure: S6 Comparison of Aminoglycoside phosphotransferase, gentamicin resistance protein gene abundance in individual samples sequenced using Illumina MiSeq or Ion Torrent platforms. (*p=0.901)

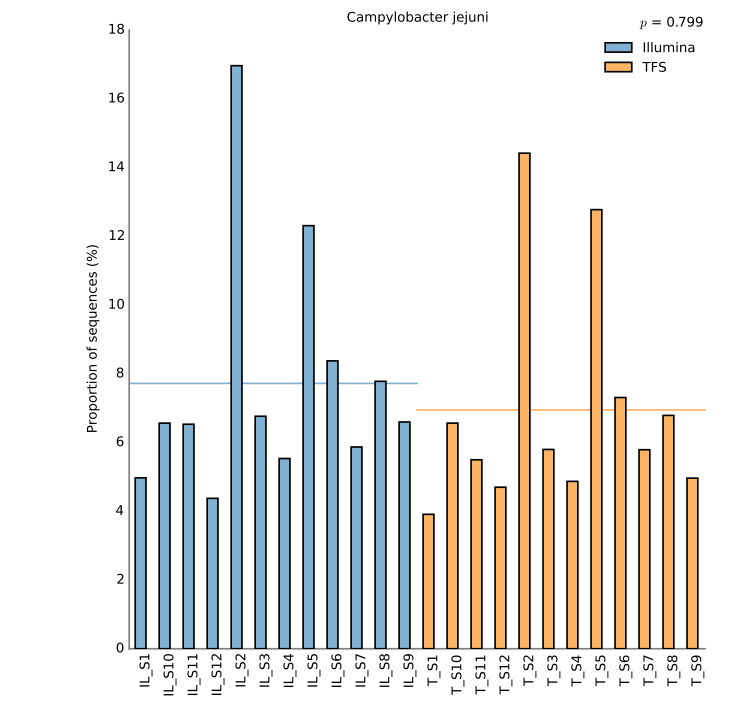

Figure: S7 Comparison of *Campylobacter jejuni* abundance in individual samples sequenced using Illumina MiSeq or Ion Torrent platforms (*p = 0.799).

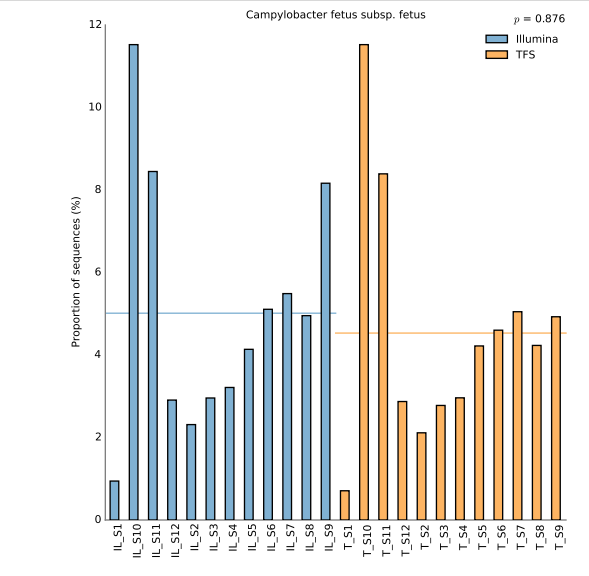

Figure: S8 Comparison of *Campylobacter fetus subsp. fetus* abundance in individual samples sequenced using Illumina MiSeq or Ion Torrent platforms (*p = 0.876).

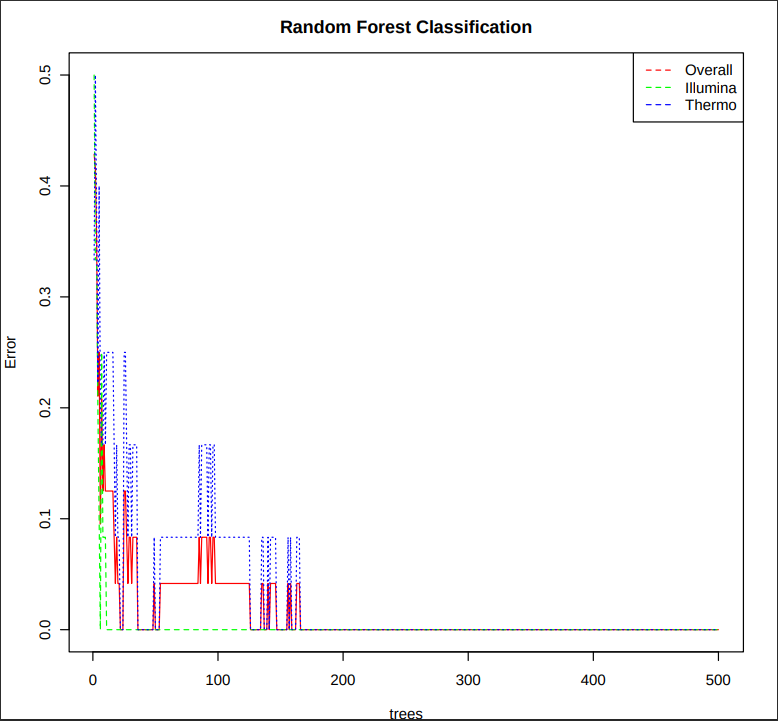

Figure: S9 Random forest classification of Illumina MiSeq vs Ion Torrent samples depicting the confidence in data analysis

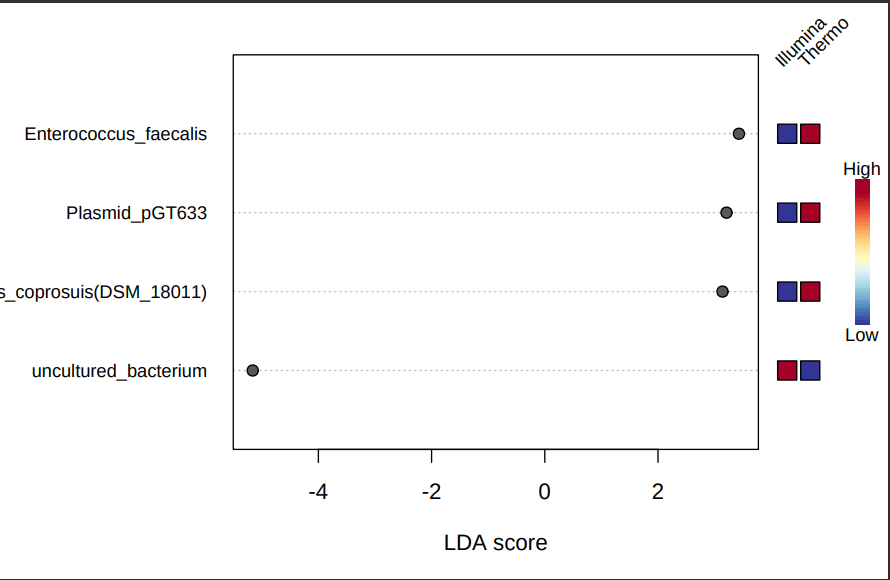

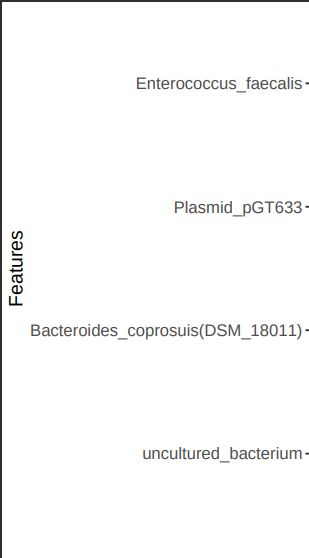

Figure: S10 LEfSe analysis for organism detection comparing Illumina MiSeq and Ion Torrent, keeping log LDA cut off as 3.0 and *p-value as 0.05.

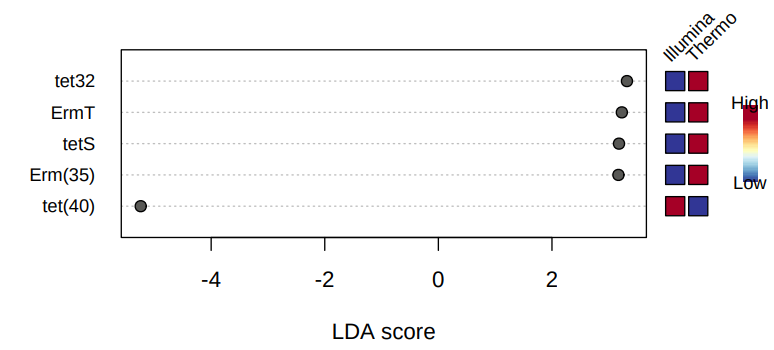

Figure: S11 The LEfSe analysis of the AMR genes keeping log LDA cut off as 3.0 and *p-value as 0.05.
